# The tobacco hornworm as a novel host for the study of bacterial virulence

**DOI:** 10.64898/2026.04.04.716455

**Authors:** Emma K. Spencer, Craig R. Miller, James J. Bull

## Abstract

The tobacco hornworm moth (*Manduca sexta*) is evaluated as a model of bacterial virulence and host-pathogen dynamics. Infections of *Pseudomonas aeruginosa* were established by injection of 5^th^-instar larvae, and multiple assays of virulence were evaluated. Infected larvae exhibited dose-dependent mortality, reduced growth, melanization, behavioral changes, and altered frass constitution. Even low-dose infections that were not fatal exhibited impaired growth, but individual growth trajectories revealed considerable heterogeneity among worms given the same dose. Twice-daily antibiotic treatment with gentamicin or cefepime improved survival four- to five-fold but did not rescue 100%. Heat-killed cells and filtered culture supernatant alone induced significant morbidity and mortality, suggesting secreted bacterial products are important to pathogenesis. Bacterial burden analysis revealed a shifting bacterial distribution over time, with decreasing hemolymph titers and increasing localization in fat body, gut, and carcass. Hornworms thus offer a more sensitive analysis of bacterial infection dynamics and consequences than do larvae of the more commonly used wax moth.

## Introduction

Bacterial infections are a leading cause of mortality worldwide, responsible for an estimated 7.7 million deaths in 2019, accounting for about one in every eight deaths globally [1, 2]. Understanding bacterial virulence and host responses requires animal models. Mammalian models, such as mice, have been instrumental [3–6], but they are limited by ethical considerations, financial cost, and the need for specialized facilities, contracts, and trained personnel [7–11]. To avoid some of the challenges of using mammals, invertebrate species have been used [12–17]. Invertebrates have a complex innate immune system and can generate highly specific innate immune system responses and thus offer a good stand-in for the more complex mammalian innate and adaptive immune system [18–23].

The common existing invertebrate models, such as *Drosophila melanogaster*, *Caenorhabditis elegans*, and wax moths (*Galleria mellonella)* are small. Their sizes restrict sampling and quantitative analyses within individuals. The result is that infection outcomes are often reduced to binary survival metrics, masking host dynamics and individual heterogeneity. Even wax moths, the largest of those three invertebrates, attain only 14 – 28 millimeters in length with an average mass of 280 mg [24, 25]. Their common and most apparent measure of bacterial virulence is percent survival, leaving many nuances of the infection inaccessible.

In this study, we evaluate the tobacco hornworm, *Manduca sexta*, as an invertebrate host for studying bacterial infections. The tobacco hornworm is a specific type of hawkmoth, a group of agricultural pests in the southern United States, feeding on plants such as tobacco, tomatoes, and eggplants. This invertebrate model has been used in research and has generated insights into insect physiology, metamorphosis, and flight mechanics [26–30]. Along with physiology, the innate immune system has been well-characterized, with 186 genes identified that have intracellular immune signaling and responses and encoding approximately 199 proteins [31]. Standardized methods are available for laboratory propagation [32]. Hawkmoth larvae attain 10 – 13 grams in their final developmental stage, the size of a small mouse [33]. This large size enables assays that are not practical with waxworms and that enable more sensitive statistics because measures are quantitative instead of binary.

The bacterial species chosen for this study was *Pseudomonas aeruginosa*. This bacterium is a gram-negative, opportunistic pathogen that causes community and hospital-acquired infections [34–36]. *P. aeruginosa* most seriously impacts critically ill and immune-compromised patients [37]. In the United States, *P. aeruginosa* causes a total of 51,000 infections in the healthcare system per year [38]. Modeling bacterial infections in a host with an innate immune system as a stand in for immunocompromised patients is an important aspect to understanding how bacterial infections are established and how they persist.

## Materials and methods

### Hawkmoth strain and preparation

The hawkmoth larvae used in this study come from a colony maintained at the University of Idaho since 2023. The colony was established from eggs supplied by Great Lakes Hornworms and have been reared from protocols described elsewhere [32]. The larvae were kept at 28 °C and 40-60% relative humidity (RH) and fed a wheat-germ-based diet supplied by Great Lakes Hornworms. All inoculations were of 5^th^ instar larvae.

### Pseudomonas aeruginosa strain and preparation

The *Pseudomonas aeruginosa* strain used in this study was PAO1, one of the most common research strains [39]. PAO1 strain was grown in Luria Broth (LB) and on LB agar plates (LB at 10 g tryptone, 5 g yeast extract, and 10 g NaCl per liter [40, 41]). Bacteria from a single colony were routinely cultured overnight in 25 mL of LB broth at 37 °C. For injections, an overnight culture was pelleted, washed in phosphate buffered saline (PBS) three times, before a final suspension in PBS to create a stock. Bacterial concentrations measured by CFUs (colony forming units) were determined by plating the same day as injections.

Heat-killed bacteria were prepared by heating a centrifuged PBS-suspended culture at 95 °C for 15 – 20 minutes, vortexed every 5 minutes. The heat-killed stock was plated to evaluate any colony growth (Supplemental Figure 1) and 10 μL of the stock was injected for ‘Heat-killed PAO1’ treatments. Filtered supernatant stocks were made from the supernatant of a PBS-washed overnight culture (15,000 g for 15 minutes) and passing it through a 0.2 μm filter. Supernatant stocks were also plated to evaluate any colony growth (Supplemental Figure 1), and 10 μL of this stock was injected for ‘Filtered Supernatant PAO1’ treatments.

### Infection methods, bacterial counts, and mass measurements

The larvae selected for bacterial virulence assays were 0 – 48 hours beyond the onset of their 5^th^ instar, with masses 1.0 – 2.5 g. Larvae were used only if robust and healthy with no atypical color phenotype. Prior to the injection, larvae were stored at 4 °C for 15 minutes, to several hours as a pseudo-anesthetization to facilitate easier handling and injection.

A 30-gauge needle and Hamilton^TM^ microliter syringe were used for injections. If a larva visibly bled hemolymph during the injection, it was discarded. Injections were done between the fourth and fifth proleg (Supplemental Figure 2), and with minimal pressure on the larva. Injection volumes of bacterial suspensions and supernatant were 10 μL. Larvae were scored for survival and weight/mass twice daily for five or more days post injection. Larval rearing conditions were kept consistent throughout all trials and rearing (with no explicit light/dark cycle at 28°C and 40 – 60% RH). All masses are reported as *relative* mass, the mass at the time of observation divided by the mass at the time of inoculation.

Tissue-specific bacterial burdens were determined by dissecting the larva into four different tissue samples: hemolymph, gut tract and fat body tissue, carcass, and collected frass. Frass was collected at the time of weighing, and previously produced frass was discarded. Hornworm larvae produce about 2 frass pellets per hour. Larvae underwent pseudo-anesthetization by placement on ice, or in the refrigerator (4 °C) for a 15-minute period prior to the dissection. The larvae, surfaces, and all tools were surface sterilized with 70% ethanol. The hemolymph was collected by either puncturing the epidermis between the first and second segment of the larva, or by cutting off the horn at the terminal segment before allowing the hemolymph to bleed into a collection tube (Supplemental Figure 3A). Dissections then proceeded by a dorsal incision and removal of the gut and attached fat body tissues, suspended in PBS (Supplemental Figure 3B). Volumetric calculations for determining colony forming units were done using the total volume of the tissues suspended in the PBS. The head was removed from the remaining tissue (carcass). The carcass was also suspended in PBS (Supplemental Figure 3CD). For frass analysis, frass was collected externally (within 15 minutes of the designated timepoint) and suspended in PBS. The gut tract and fat body, carcass, and frass samples were all homogenized separately in PBS with a sterile Tissue-Tearor^TM^, then vortexed for ten seconds. Dilutions were plated on LB, MacConkey, and Cetrimide agar plates to determine CFUs.

### Hemolymph draws

We sought a way to extract only a small volume of hemolymph that subjected the worm to no more than a mild impact on growth. A sterile, size 15 beadsmith needle was used to penetrate the skin of the larvae in the terminal segment; pressure was applied to the abdomen to facilitate bleeding. A pipette tip was used to collect the hemolymph that beaded on the worm surface, typically a volume of 20 – 50 μL. (A 30-gauge needle was, by contrast, too large; the volume of hemolymph extruded often exceeded 100 μL, and bleeding was slow to stop).

### Selective media to differentiate background microbiota from *Psuedomonas*

The microbiome of the tobacco hornworm larvae is considered to be transient, not residential and thus likely originating from external sources [42, 43]. Bacterial species present in the colony of hornworms were established to be mostly gram-positive cocci species [32]. Selective media was used to differentiate between *P. aeruginosa,* and the bacteria present in the microbiome: Cetrimide [44] and MacConkey agar [45]. When plated on MacConkey agar, *P. aeruginosa* colony morphology is round, flat, and colorless, as it is a lactose non-fermenter [46]. When plated on Cetrimide agar, *P. aeruginosa* colonies are a fluorescent yellow green [44].

### Antibiotic administration

Cefepime and gentamicin were the two antibiotics used in this study to treat experimental infections. Cefepime is a beta-lactam, inhibiting cell wall synthesis, leading to autolysis and bacterial death [47]. Gentamicin is an aminoglycoside and inhibits bacterial protein synthesis by binding to the 30s ribosome [48].

Cefepime and gentamicin treatments were administered four to six hours after the beginning of the bacterial infection and were given twice daily for five days. Antibiotic stocks were made to a concentration of 5 mg/mL in 0.9% saline and were diluted so that the 10 uL dose was from a concentration of 50 μg/mL (thus a worm received 0.5 μg at each inoculation). Antibiotic treatments were injected between the first and second set of prolegs. Control treatments consisted of *Pseudomonas*-free hornworms given antibiotics twice daily for five days.

### Statistical analysis

All statistical analyses were performed in R Studio [49]. Percent survival, relative change of mass, and bacterial burden were plotted using ggplot2 [50]. For percent survival analysis, Kaplan-Meier followed by a log-rank (Mantel-Cox) test using the Survival package was used. A Cox proportional hazards model was used to quantify effect of dose-dependency on death over time, using the coxph() function in the survival package [51]. Model significance was evaluated using the likelihood ratio test, and proportionality of hazards was assumed.

To evaluate the effects of a PAO1 dose on larval growth, statistical analyses incorporate both longitudinal and end-point based approaches. Larval relative mass data were analyzed using linear mixed-effects models to evaluate the differences in growth trajectories across bacterial dose treatment, and to assess variance structure over time. The models included treatment group, day post-injection (DPI), and their interaction as fixed effects. Individual larvae were modeled as random effects to account for repeated measures, and fixed effects included day post-injection and its interaction with treatment [52].

In addition, endpoint relative growth was evaluated using a stepwise analysis of variance (ANOVA) framework to test treatment effects at the population level. An omnibus ANOVA with treatment group as fixed effect was first used to assess overall differences in endpoint relative growth among groups. Upon detection of significant main effect, post hoc pairwise comparisons were conducted using estimated marginal means with Tukey’s adjustment for multiple comparisons, as implemented in the emmeans package [53].

Inter-individual variation in larval growth was quantified using endpoint relative growth among surviving larvae. For each treatment group, growth heterogeneity was summarized as the standard deviation of relative growth values among surviving individuals at day 4 post-injection. Groups with one or no surviving larvae at this time point were excluded from variance calculations.

For sequential samples, repeated hemolymph was obtained from the same individuals and sample size was limited, statistical analyses focused on biologically meaningful contrasts rather than timepoint-specific comparisons. Differences in total bacterial burden between surviving and non-surviving larvae were assessed using a Wilcoxon rank-sum test [54]. Effect size was quantified using rank-biserial correlation to evaluate the magnitude of the association between bacterial burden and host mortality [55, 56].

To quantify bacterial dynamics during infection, bacterial concentrations were measured over time in multiple compartments: hemolymph, fat body and gut, carcass, and excreted frass. For each larva, total internal bacterial burden was calculated by summing bacterial counts across all internal tissues at each sampling time point. Temporal changes in total internal bacterial concentration were analyzed using a generalized linear model (GLM) and Gaussian error distribution with log link function, with time post-injection treated as a continuous predictor [57]. A separate GLM with a log link was fitted to frass-associated bacterial concentration to evaluate temporal trends in bacterial excretion. Model coefficients, standard errors, and associated t-statistics were used to assess the effect of time on bacterial concentrations.

## Results

### Dose-dependent mortality

Groups of 10 to 25 fifth instar larvae were infected with increasing doses of PAO1. Infected larvae were held at 28 °C and were scored daily for survival and weighed for five consecutive days. Larvae died in a dose-dependent manner (Fig 1): the more bacteria injected, the higher the mortality, and at a faster rate (χ^2^ = 32.0, df = 5, p = 6 x 10^-6^). A log-rank (Mantel-Cox) test revealed significant differences in survival among bacterial dose groups (χ^2^ = 18.7, df = 7, p = 0.009). Specifically, higher doses of PAO1 resulted in greater and more rapid larval mortality. Cox proportional hazards modeling showed that bacterial dose significantly predicted larval survival (likelihood ratio test: χ^2^ = 22.61, df = 7, p = 0.002).

**Fig 1.**
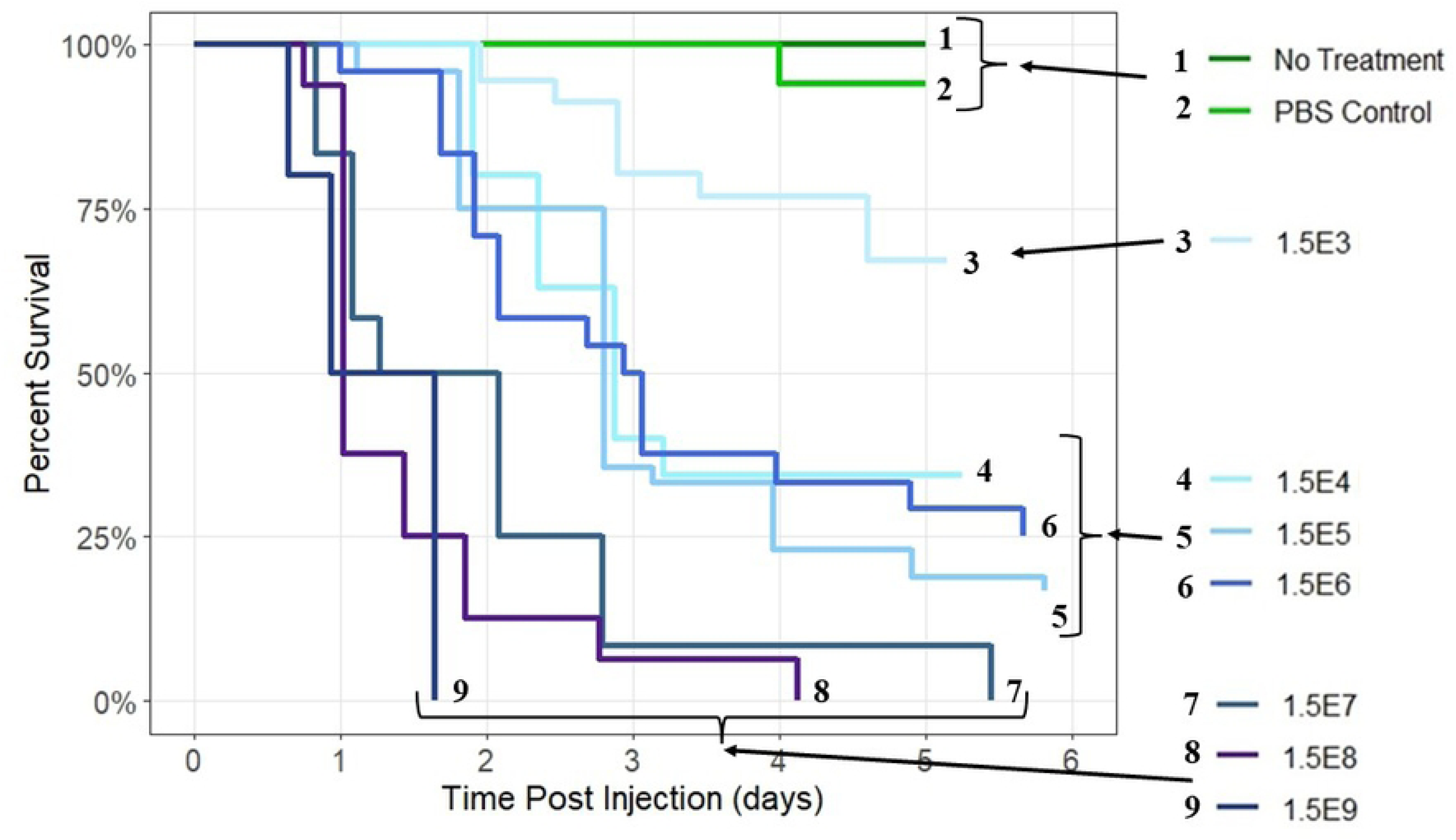
Percent survival curves for larval groups injected with increasing doses of *P. aeruginosa* (PAO1). All groups had n ≥ 8 individuals (No Treatment Control N = 8, PBS Control N = 36, 1.5E3 N = 25, 1.5E4 N = 35, 1.5E5 N = 34, 1.5E6 N = 24, 1.5E7 N = 12, 1.5E8 N = 16, 1.5E9 N = 10). The two controls are ‘No Treatment Control’ and ‘PBS Control’, which respectively are larvae that had no injection and larvae that were injected with only phosphate buffer. Survival of these two control groups were 100% and 94%, respectively. Injection of bacteria resulted in decreased survival, more so with higher doses. The lowest dose (1.5E3 CFU) experienced 67% survival, the highest dose (1.5E9 CFU) experienced 0% survival in less than 2 days.

The two controls (No Treatment Control and PBS Control) were combined as their survival didn’t differ between them (χ^2^ = 0.5, df = 7, p = 0.5). Pairwise Cox proportional hazards comparisons using the pooled control groups revealed that all PAO1 doses significantly increased larval mortality risk relative to the controls (Holm-adjusted p < 0.01 for all doses). Hazard ratios increased monotonically with bacterial dose, ranging from 7.9-fold higher risk at 1.5E3 CFU to over 350-fold higher risk at 1.5E9 CFU (Table A1). These results also confirm that hornworms are susceptible to PAO1 in a dose-dependent manner.

### Dose-dependent growth (morbidity)

The large size of fifth instar hornworms and their rapid growth from approximately 1 to 10 gm allows for a sensitive assay of growth. The same larvae studied for survival were weighed daily while alive or through day 6 post-injection. Growth shows a more detailed picture of the infection than survival, and indeed, is a more sensitive measure of the bacterial infection. Growth here is always presented as *relative* growth, calculated as an individual’s weight divided by its starting weight at the time of injection (t = 0). A relative growth value above 1 indicates weight gain, whereas values below 1 indicate loss of mass, interpreted as morbidity. We also observed that larvae often declined in weight before dying. Larvae that died were assigned a weight of zero at the time of death to contrast from survivors. All deaths were omitted from the average growth curves shown in Figure 2.

**Fig 2.**
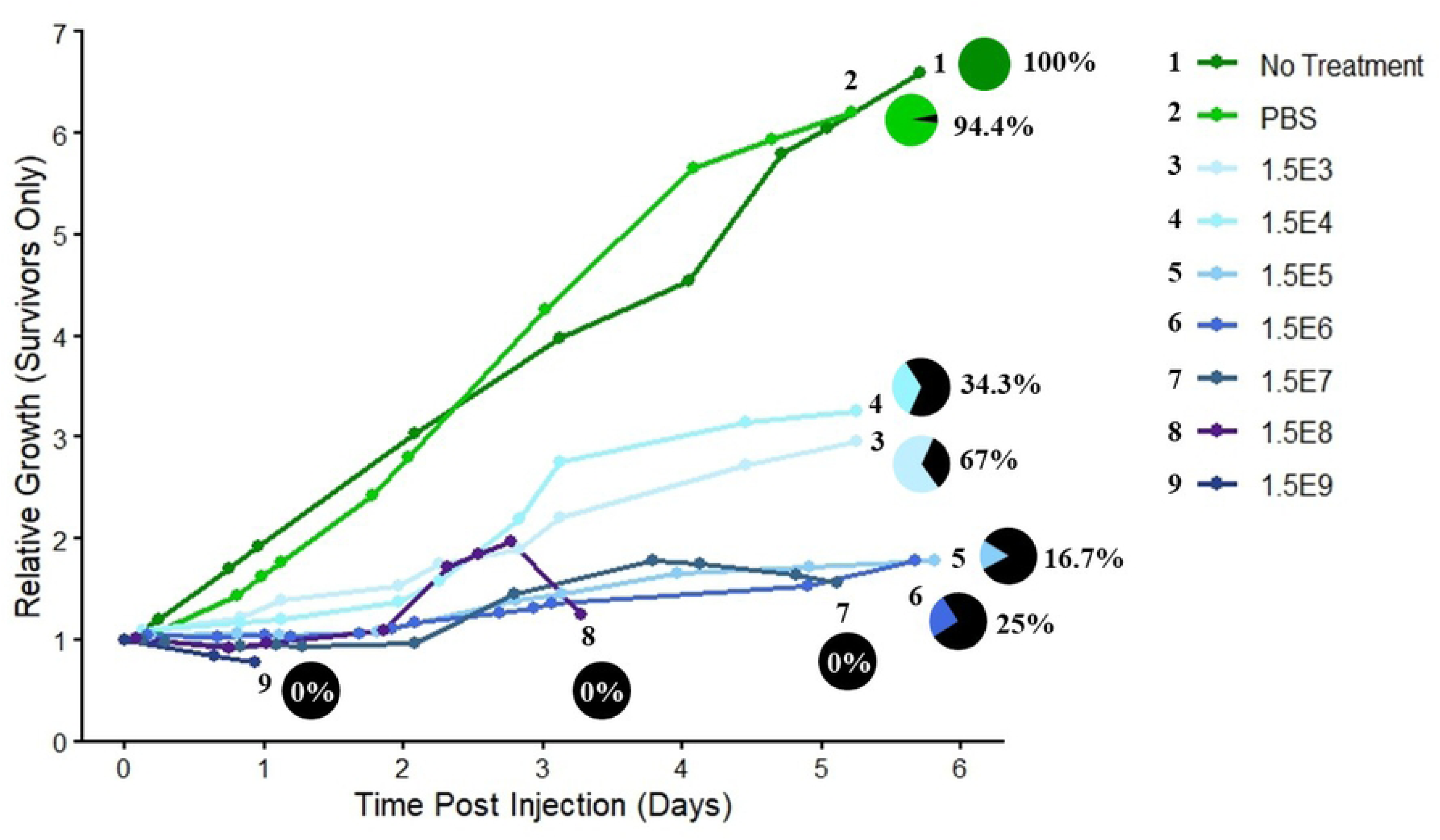
Mean relative growth of surviving larvae following the injection of different doses of PAO1. Growth values represent averages conditional on survival at each time point; deceased individuals were excluded from the averages. The growth curve terminates when no surviving larvae remains. The terminal percent survival at 5-6 days (Figure 1) is given by the pie charts. All groups had N = 8 or higher (No Treatment Control N = 8, PBS Control N = 36, 1.5E3 N = 24, 1.5E4 N = 35, 1.5E5 N = 34, 1.5E6 N = 24, 1.5E7 N = 12, 1.5E8 N = 16, 1.5E9 N = 10). All groups infected with PAO1 had significantly reduced growth rates compared to the controls (No Treatment & PBS Control) (F_5, 29.5_ = 119.5, p < 0.001). The two lowest doses (1.5E3 and 1.5E4) did not differ from each other (p = 0.894), whereas the higher doses (1.5E5 and above) caused markedly greater suppression and differed significantly from the lower doses (p ≤ 0.0187). No difference was detected once again between the higher doses, suggesting a plateau in growth inhibition above the 1.5E5 threshold.

Larval relative growth trajectories were analyzed using a linear mixed model that incorporated repeated measurements across time to evaluate dose dependency. Growth differed significantly among groups (LMM: F_5, 29.5_ = 119.5, p < 0.001), with differences evident by day 4 post-injection. The PAO1-infected groups exhibited significantly reduced growth relative to both uninfected controls (No Treatment and PBS). Among infected larvae, relative growth declined with increasing dose. The highest dose in which at least some worms survived 5-6 days (and thus could be used for measures of final growth) was 1.5E6 CFU, and this dose exhibited significantly reduced growth compared to the two lowest doses (1.5E3 and 1.5E4).

To complement trajectory-based analyses, we also examined endpoint growth at day 4 for individual larvae, providing a direct assessment of dose-dependent heterogeneity in growth outcomes. One-way ANOVA of day 4 relative growth revealed significant differences among all groups (F_5, 78_ = 54.96, p < 0.001) (Table A2). This heterogeneity remained significant after removal of uninfected control groups (No Treatment and PBS) (F_3, 38_ = 6.02, p = 0.0019), indicating persistent variation in growth outcomes among infected larvae across PAO1 doses. However, when the lowest PAO1 doses (1.5E3 and 1.5E4 CFU) were excluded, no significant heterogeneity remained among higher doses (F_1, 15_ = 0.97, p = 0.34) (Table A3).

### Individual variation in growth

Figure 2 presented the average growth rate of the larvae that survived the infection. Figure 3 illustrates individual growth trajectories for each group, highlighting the inter-individual heterogeneity in the growth in response to the infection. At Day 4 post-injection, the growth heterogeneity was quantified as the standard deviation in relative growth among surviving larvae. Control groups exhibited modest variation in growth (No Treatment: SD = 0.70, PBS: SD = 0.89), consistent with normal growth we have observed in other contexts (32).

**Fig 3.**
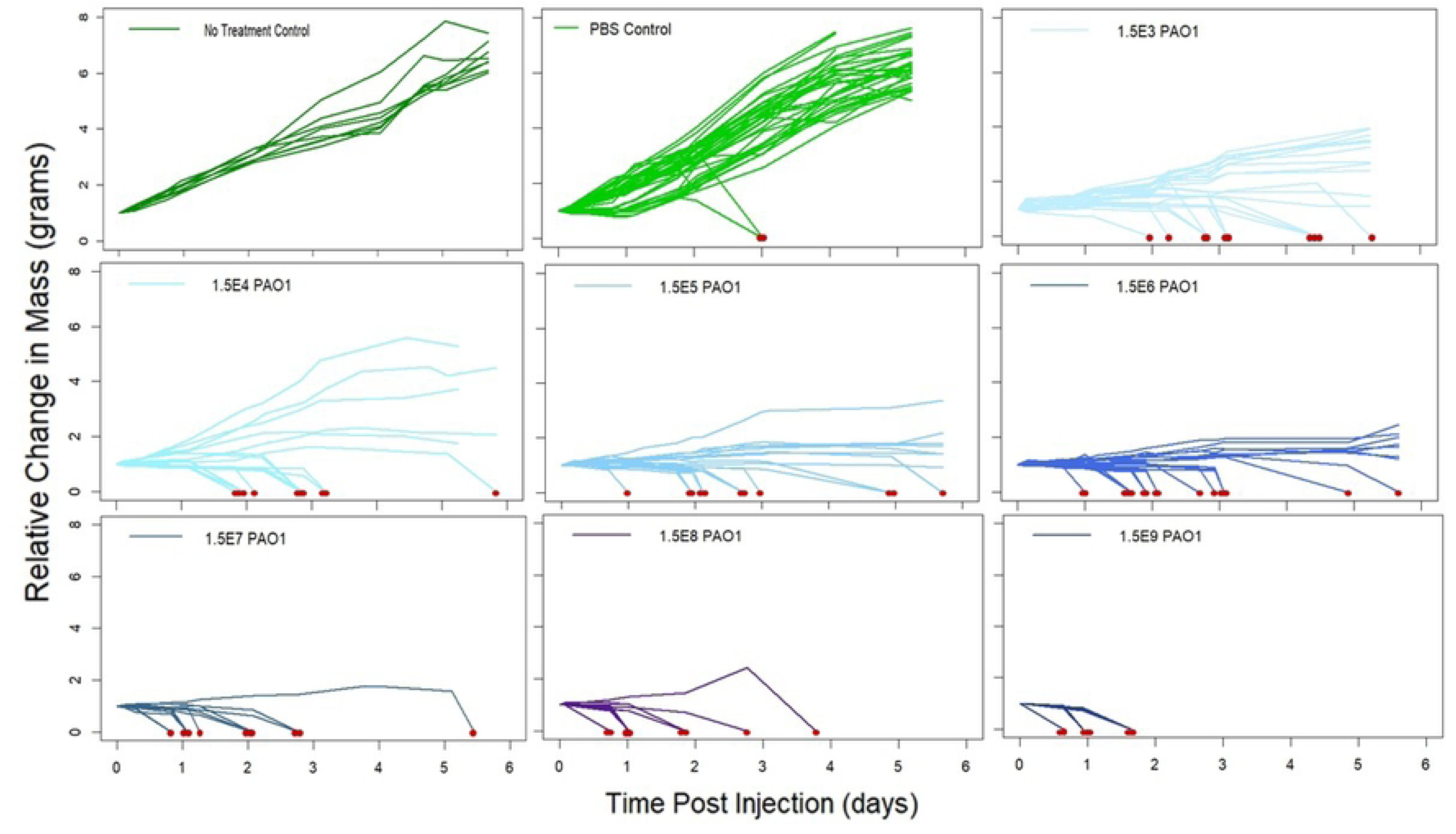
Individual growth trajectories for the treatment groups of Figure 1 and 2. Each line represents an individual larva, and the red dots indicate the death of the individual. All groups had N = 8 or higher (No Treatment Control N = 8, PBS Control N = 36, 1.5E3 N = 25, 1.5E4 N = 35, 1.5E5 N = 34, 1.5E6 N = 24, 1.5E7 N = 12, 1.5E8 N = 16, 1.5E9 N = 10). The two controls (‘No Treatment Control’ and ‘PBS Control’) were not injected with bacteria. Though there is some variability with the PBS Control group, the controls exhibited far less variability in growth across time than do the groups receiving bacteria. Additionally, larvae that survived an infection typically did not grow at the same rate as individuals in the control groups, except for some individuals in the 1.5E4 dosage group. In doses 1.5E3 through 1.5E6, there is a lot of variability in individual growth rates. For doses 1.5E7 and 1.5E8, there is less variability because growth is so drastically suppressed. At 1.5E9, too few individuals survived even a few days to warrant statistical analysis.

In contrast, the low PAO1 doses (1.5E3 – 1.5E4 CFU) displayed pronounced heterogeneity in growth outcomes among the survivors, with variability increasing from 1.5E3 (SD = 0.95) to a peak at 1.5E4 (SD = 1.43). At intermediate doses (1.5E5 – 1.5E6 CFU), both mean growth and heterogeneity declined sharply, reflecting uniform growth arrest among the surviving larvae (SD < 0.55). At higher doses (≥1.5E7 CFU), rapid mortality resulted in one or zero surviving larvae by day 4, preventing meaningful assessment. In comparison to Figure 1, Figures 2 and 3 show more nuanced information regarding the infection consequences. With these measures, it is possible to evaluate mortality and morbidity (Table A4).

### The effect of sequential sampling on growth

The large size of hornworms raises the prospect of sequentially sampling hemolymph to measure bacterial concentrations. But does removal of hemolymph affect growth or survival? Seven uninfected individuals were subjected to hemolymph drawn five times over the course of four days and their relative growth and volume of hemolymph drawn were recorded. The relative growth of these larvae was significantly less than the growth of the ‘No Treatment Control individuals’ (F = 8.78, p = 0.0004; Fig 4), although the magnitude of growth reduction is modest. The volume of hemolymph draws was between 15 – 100 μL with the average draw about 40 μL. At time 0, this volume represents about 2.5 – 3% of the larva’s body mass, while at day 4, this is only about 0.7% of the total body mass. The total of hemolymph drawn from the larvae over the five draws was between 150 – 250 μL. Individual growth rates declined with increasing cumulative hemolymph removal, but this relationship was not statistically significant.

**Fig 4.**
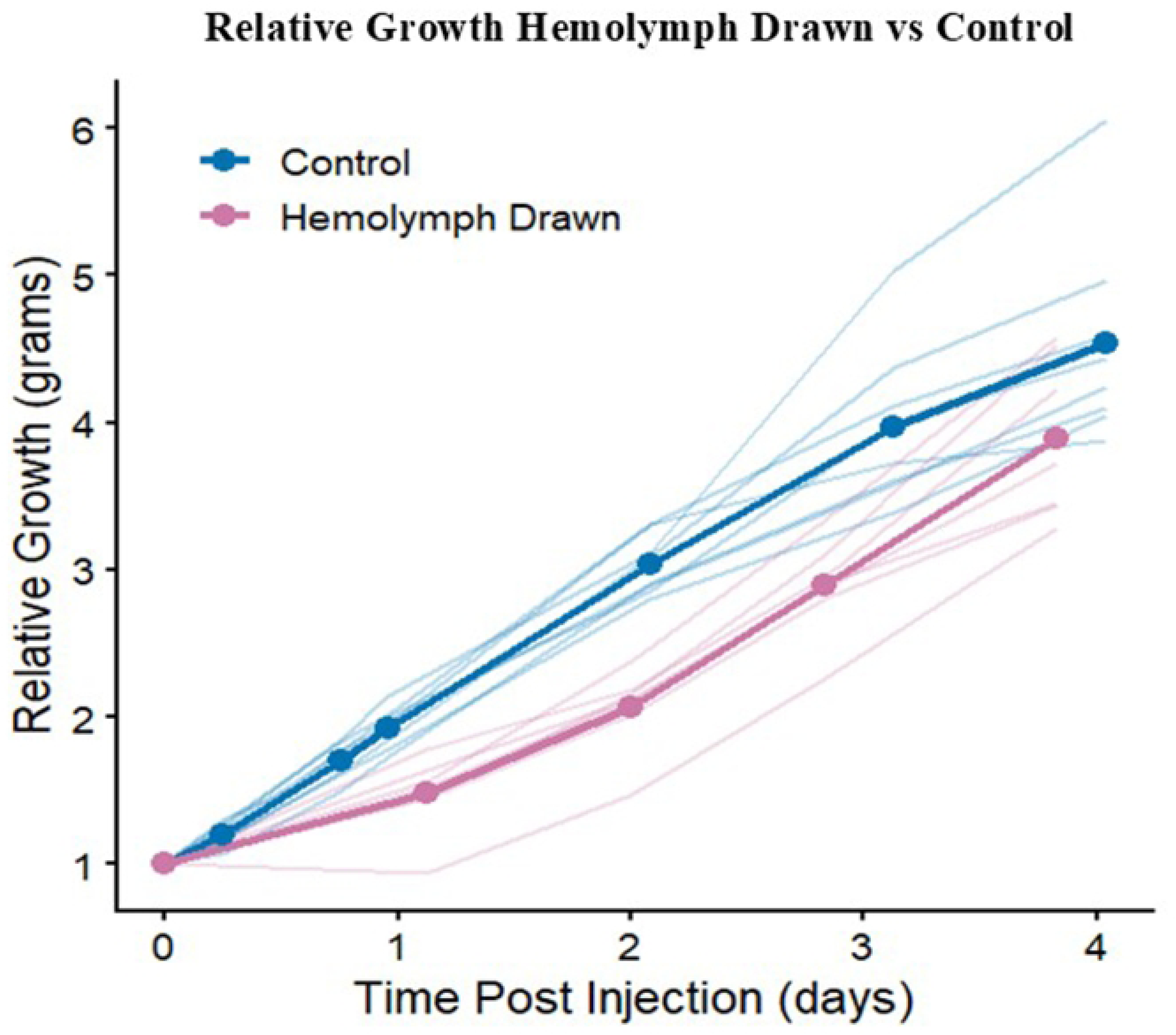
Relative growth of larvae of hemolymph drawn compared to controls. Seven uninfected larvae had hemolymph drawn five times over the course of four days. The relative growth of the hemolymph drawn larvae compared to the ‘No Treatment Control’ individuals (Figure 3). The transparent lines represent individual larvae, the opaque lines represent group averages. The hemolymph-drawn group has decreased growth compared to the unmanipulated group (F = 8.78, p = 0.004), but the effect is modest.

### Sequential sampling with bacteria

To determine bacterial counts in hemolymph over time with sequential draws, approximately 2E5 PAO1 were injected into each of eight larvae and their hemolymph was drawn once per day over four days. Weight, bacterial counts and the volume of hemolymph removed were recorded (Fig 5).

**Fig 5.**
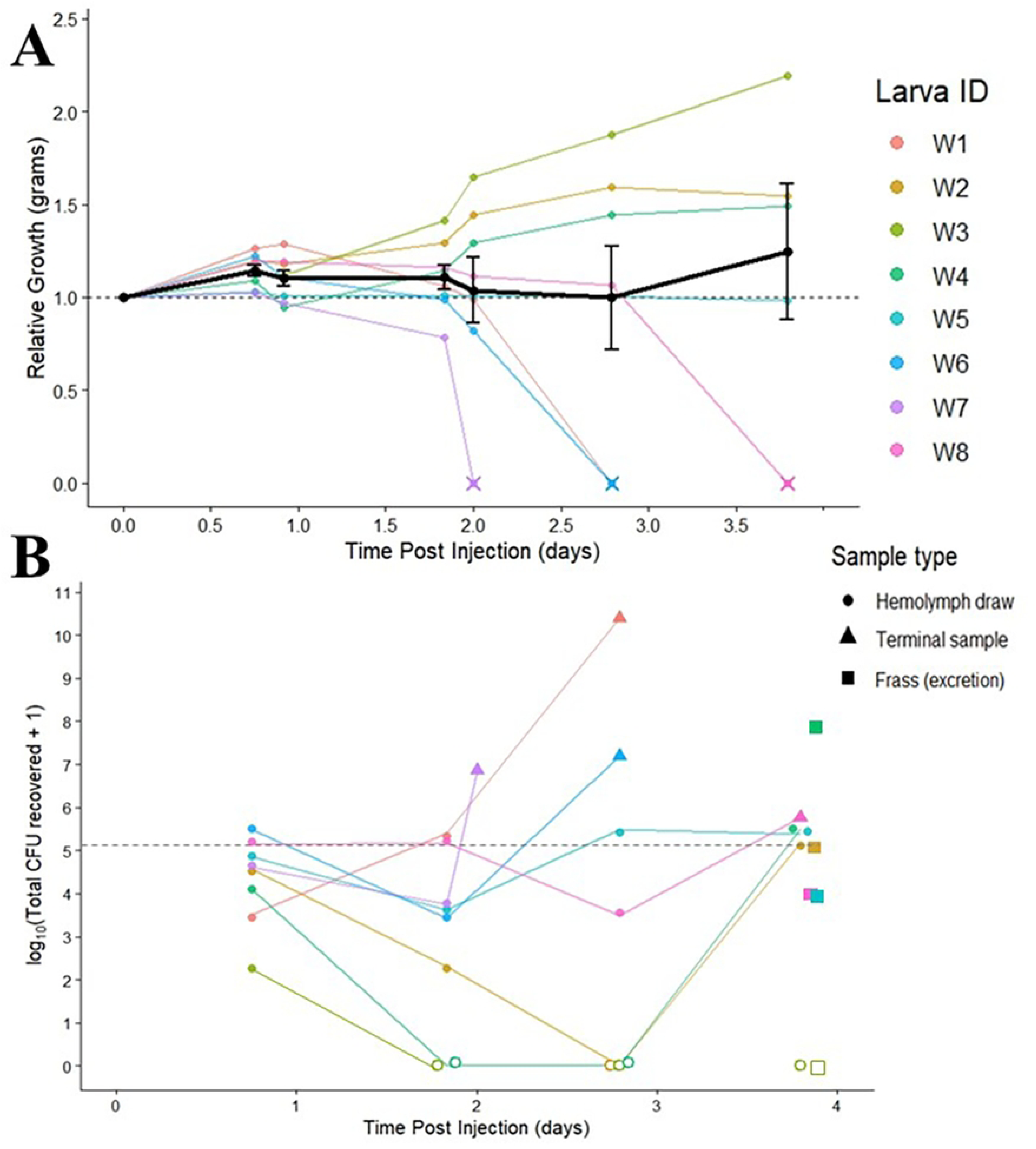
Hemolymph sequential sampling of larvae infected with bacteria. Eight individual hornworms were injected with 2E5 CFU of PAO1 at time 0 and monitored over four days. Hemolymph was collected once daily (or until death), and the bacterial concentrations were quantified for each sample. Four of the eight larvae died between 0 and 3.8 DPI (W7: 2 DPI; W1 and W6: 2.8 DPI; W8: 3.8 DPI). Three additional larvae died between 4 – 5 DPI (not shown), and only one larva (W3) survived beyond Day 5 (12.5% survival). (A) Individual growth trajectories. Larvae that died were assigned a relative growth of 0 at the time of death; the black line shows the mean of surviving larvae ± SD. (B) Bacterial concentrations over time. Filled circles indicate samples from live larvae, open circles denote no detectable colonies (0 CFU in 10 μL), triangles represent terminal samples collected at death (hemolymph, gut, and fat body combined), and squares indicate bacterial concentrations in the frass collected at the final sampling point (per gram).

The relative growth trajectories of surviving larvae for DPI 0 – 4 did not differ between the group injected with 1.5E5 and 2E5 cells (LME: p ≥ 0.73, Welch’s t-test: p = 0.53). Considerable heterogeneity is observed in bacterial counts of the same worm over time. Except possibly for the last sample taken before death, there is no obvious increase in bacterial density.

### Whole-body bacterial loads over time

To assess the dynamics and spatial analysis of the bacteria during the infection, bacterial counts were measured in different tissues over time, including frass (Fig 6, Fig S2). There was no statistically detectable increase in total internal bacterial concentration over time, consistent with the results from sequential hemolymph sampling. A generalized linear model (GLM) with a log link function was fitted to total internal bacterial concentration (summed across all internal tissues per larva) as a function of time. The effect of time was not statistically significant (β = 0.5446 ± 0.4019, t (23) = 1.355, p = 0.189), indicating no significant net change in internal bacterial burden over the sampling period (Figure 6C). In a separate model, the frass-associated bacterial concentration showed a borderline-significant decrease over time approached, with a negative slope (β = -0.3296 ± 0.1629, t (20) = -2.024, p = 0.057) (Fig 6C). We of course cannot reject a model of bacterial growth in the host, but we have no evidence of it from these data.

**Fig 6.**
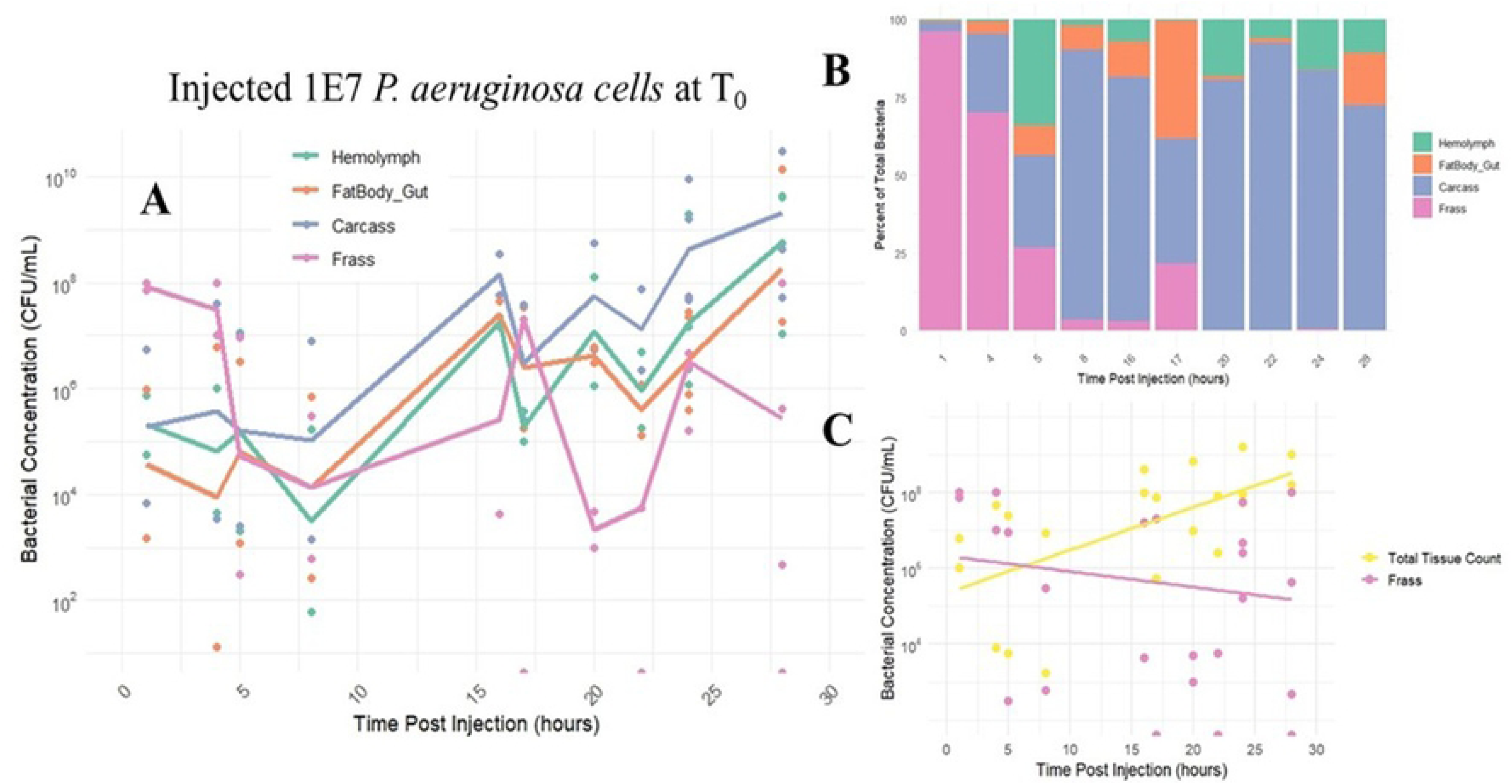
Distribution, location, and overall number of PAO1 cells present in individual hornworms over time in different tissues. A) The cell concentration (CFU/mL) over time in each of the different tissue types: hemolymph, fat body & gut, carcass, and excreted frass. Each data point indicates the concentration of bacteria observed at the time point. There is a high degree of variability within all tissue types, but there is a trend of increasing cell count in the carcass, fat body, and hemolymph over time. B) The proportion of the total bacteria observed at the time point split into the tissue type. After the first few hours, the proportion of the bacteria seen is dominated by the carcass. C) The number of bacteria for the hemolymph, fat body and gut, and carcass were all summed together and plotted against bacteria in the frass. A generalized linear model with a log link was fit to assess the change in total internal bacterial concentration over time. The effect of time was not statistically significant (β = 0.5446 ± 0.4019, t (23) = 1.355, p = 0.189), indicating no significant change in bacterial load. A similar model was fitted for frass-associated bacterial concentration. The effect of time approached significance, with a marginal negative slope (β = -0.3296 ± 0.1629, t (20) = -2.024, p = 0.057).

As bacteria were injected directly into the hemocoel, circulating hemolymph was expected to harbor higher bacterial concentrations than other tissues. Surprisingly, initial samples revealed that a substantial proportion of the bacteria was already located outside of the hemolymph, with the highest early concentrations observed in the frass (Fig 6B). This redistribution is consistent with the sequential hemolymph sampling experiments, which showed declining hemolymph bacterial loads despite continued infection, and suggests that the bacterial loss from circulation occurs rapidly following the injection. However, individual larvae exhibited substantial variability in tissue-specific bacterial burdens, indicating that these dynamics are not uniform.

These observations suggest the intriguing possibility that bacterial transfer into frass may help clear the infection or at least slow the increase of bacterial density within the host. If the actual number of bacteria being transferred into the frass was known, it would be possible to calculate a minimum within-host bacterial growth rate by accounting for the loss to the frass. However, it is not straightforward to determine the actual number of bacteria transferred to frass from the bacterial counts observed in the frass because frass provides nutrients for bacterial growth (Fig S4). The counts observed in frass are undoubtedly larger than the number of bacteria transferred to frass due to bacterial growth in the frass before and after excretion from the worm.

Larvae will consume their frass if it is not removed from the containers, creating a potential route for re-exposure to excreted bacteria. However, oral gavage with PAO1 had no detectable effect on larval growth or survival. This suggests that re-ingestion of the bacteria via frass is unlikely to substantially influence infection outcomes, although the extent of this process remains uncertain.

### Bacterial growth in dead hornworms

To determine the role the immune response might play in the infection and potential replication of PAO1, the concentration of bacteria was evaluated in dead larvae. Eight fifth instar larvae were put in the freezer (-20 °C) for 24 hours and were then brought back to room temperature (22 °C) before approximately 1E7 cells were injected into the dead worms, then kept at 28 °C. Larvae were then ground up and plated on selective agar to determine the concentration of *Pseudomonas* over time. Two individuals were destructively sampled at each of 2, 4, 8, and 17 hours post injection (Fig 7). In the first four hours, there was no detectable change in the concentration of PAO1. However, at hours 8 and 17, the concentration had increased by 2 orders of magnitude. This increase in concentration was not observed in the live hornworms (Fig 6), which suggests that the immune system may be suppressing bacterial replication

**Fig 7.**
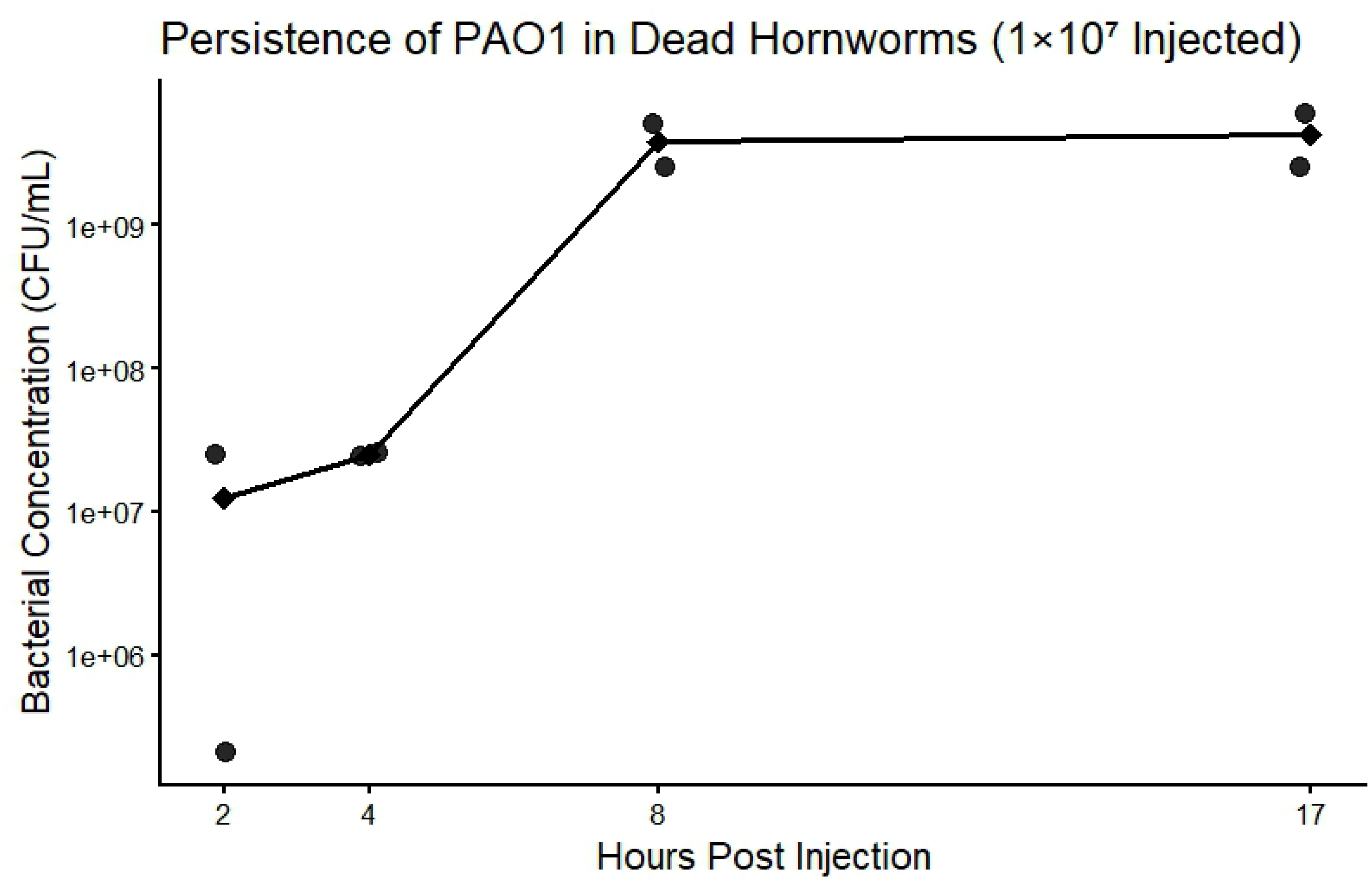
Persistence and replication of PAO1 in dead hornworm larvae. 1E7 PAO1 was injected into eight dead larvae, and the concentration of bacteria was evaluated at 2, 4, 8, and 17 hours post injection.

### Antibiotics rescue but only partly

Survival and relative growth were recorded following bacterial infection and antibiotic treatment (Fig 8). Larvae were injected with 2.5E6 cells and treated with two types of antibiotics twice daily for five days. Two additional groups received only antibiotics for the same frequency and duration as controls. The antibiotics themselves had no detectable effect on survival, with 93% survival in the gentamicin control, and 100% survival in the cefepime control.

**Fig 8.**
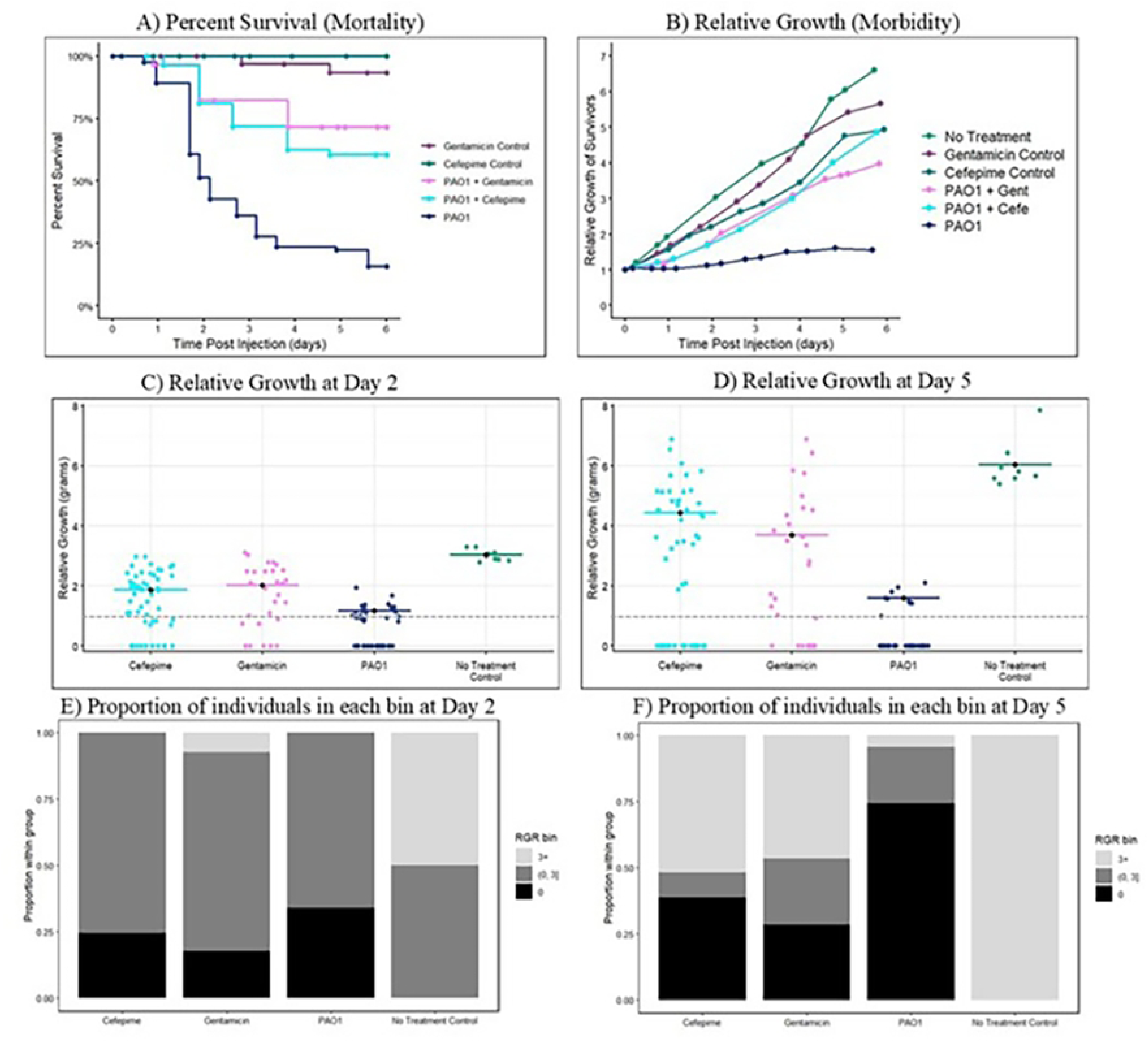
Percent survival and relative growth rates of larvae that are treated with two types of antibiotics. The groups are: No Treatment (n = 8), Gentamicin Control (n = 29), Cefepime Control (n = 30), PAO1 + Gentamicin (n = 28), PAO1 + Cefepime (n = 53), PAO1 (n = 45). A. Survival of larvae injected with 2.5E6 PAO1 cells and treated twice daily with gentamicin or cefepime for five days. Antibiotic treatment significantly increased survival relative to untreated infection (gentamicin: p = 3.3E-6, cefepime: p = 8.74E-7). B. Relative growth of surviving larvae over time. Growth trajectories differed significantly among treatment groups (p < 0.0001), and day 4 (endpoint) relative growth was significantly reduced in PAO1-infected larvae compared to the controls (p < 0.0001). Antibiotic treatment partially mitigated infection induced growth suppression but did not restore growth to the uninfected control levels. C. The relative growth of each individual larvae at Day 2 (4^th^ dose of antibiotics). Dead larvae are represented by zeros but are not included in the group growth mean (line). Dotted line is at 1 to show growth. D. The relative growth of each individual larvae at Day 5 (8^th^ dose of antibiotics). Dotted line is at 1 to show growth. Dead larvae are represented by zeros but are not included in the group growth mean (line). E & F. Distribution of relative growth for individual larvae for each group on Day 2 (E) and Day 5 (F) for each group. For ease of visual comparison, relative growth rates have been binned into 3 categories of no growth (0), between 0 and 3-fold increase, and greater than 3-fold increase.

Antibiotic treatment substantially improved the survival following the infection, although the survival remained below that of the bacteria-free controls. Only 15.7% of larvae survived the untreated infection over six days, whereas survival increased to 71.4% with gentamicin and 60.4% with cefepime (Fig 8A). Thus, antibiotic-treated larvae were four-to-five-fold more likely to survive infection than untreated larvae (gentamicin: p = 3.3E-6, cefepime: p = 8.7E-7).

Antibiotic treatment also improved larval growth. Growth trajectory analyses showed significant treatment-specific differences over time (F_5, 1454_ = 57.38, p < 0.0001). As expected, infection with PAO1 caused a pronounced reduction in growth relative to all control groups (p < 0.0001). Treatment with either antibiotic significantly increased growth relative to the untreated infection (p ≤ 0.0006). However, growth in antibiotic-treated larvae remained significantly lower than in uninfected controls (one-way ANOVA, p < 0.0001). Consistent with this pattern, the control groups (No Treatment and both antibiotic-only controls) showed similarly high growth and did not differ significantly from one another p > 0.5; Table A5).

Together, these results indicate that antibiotic treatment substantially improved both survival and growth following infection but did not fully restore larvae to the uninfected state.

### Inactivated bacteria and supernatants are harmful

An unexpected finding was the observation of mortality and morbidity from inactivated and washed, heat-killed *P. aeruginosa* cells, as well as from filtered bacterial supernatant. A variety of treatments were attempted: live cells, dead cells, supernatants from live cells, as well as with and without antibiotics (Fig 9). There were deleterious effects on survival and/or growth from all derivatives of *Pseudomonas*, and as expected, antibiotics provided no benefit except against live cells. *P. aeruginosa* is known to secrete toxins and enzymes that contribute to pathogenesis [58]. Previous studies have demonstrated that bacterial culture supernatants can retain biologically active compounds, which may impact the host physiology or survival [59–64]. Our findings that washed dead cells and supernatant could be lethal are thus not anomalous in the context of other studies. Heat-killed cells were also recognized to reduce survival in waxworms (*G. mellonella*) [65, 66].

**Fig 9.**
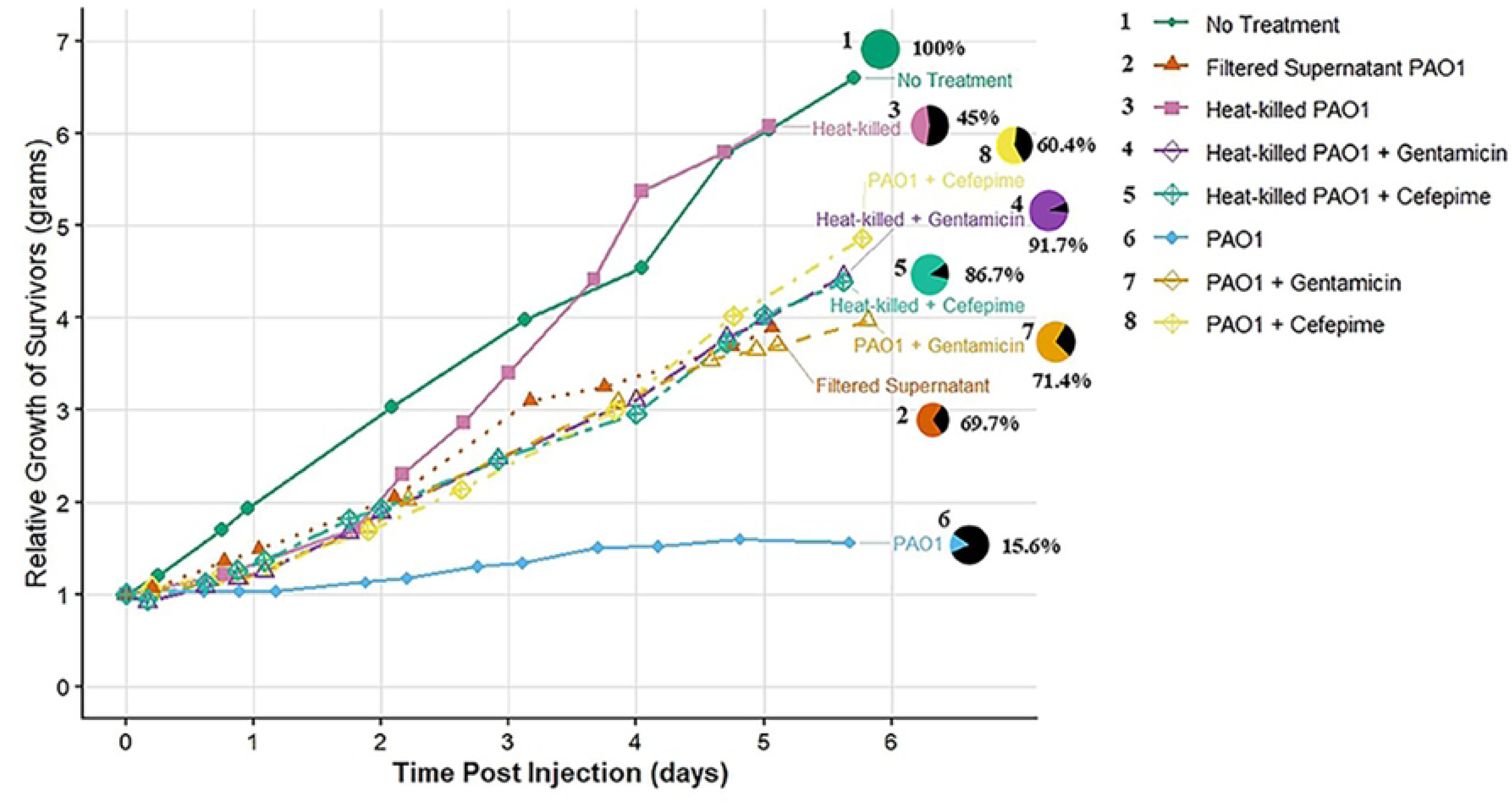
Survival and average growth of surviving larvae following exposure to live PAO1, bacterial products, and antibiotic treatments. Relative growth rates (population averages) of surviving larvae are shown over time for each treatment group. Pie charts indicate final percent survival for each treatment. PAO1 live cells caused severe growth suppression and the lowest survival. Exposure to filtered supernatant or heat-killed PAO1 resulted in intermediate survival and reduced growth relative to no-treatment control, indicating sublethal morbidity in the absence of live-bacterial replication. Antibiotic treatment during live bacterial infection substantially improved both survival and growth, while heat-killed PAO1 combined with antibiotics produced the highest survival and near-control growth. Overall, treatments differed significantly in both survival (p < 0.001), and growth rates (p < 0.001).

Live PAO1 cells resulted in the lowest survival, with only 15.7% of the larvae surviving to the end of the observation period. Larvae given heat-killed PAO1 had intermediate survival at 45%. The next groups experienced similar survival rates: Filtered supernatant at 69.7%, antibiotic treatment of live cells (PAO1 + gentamicin: 71.4%; PAO1 + cefepime: 60.4%), and antibiotic treatment of heat-killed cells (heat-killed + gentamicin: 91.7%; heat-killed + cefepime: 86.7%), and the No Treatment control at 100% (Statistics in Table A6; we note here, however, the surprising result that the greatest reduction in mortality risk was heat-killed cells and antibiotics)

Expectedly, along with lowest survival, the live PAO1 cells resulted in the lowest growth rate of all treatments. Notably, survival outcomes and growth response were not strictly coupled across treatments. For example, heat-killed PAO1 exposure reduced survival to 45%, yet the growth of surviving larvae did not differ significantly from the corresponding antibiotic-treated heat-killed groups. Conversely, antibiotic treatment nearly doubled the survival in heat-killed exposures but did not significantly increase endpoint growth relative to heat-killed alone (Statistics in Table A7). These results indicate that distinct bacterial components and treatments differentially affect mortality and morbidity, and that survival alone does not fully capture the physiological impact of exposure to *P. aeruginosa* or the byproducts it produces.

### Qualitative Results

Larvae exhibiting symptoms from the bacterial infection underwent a reduction in body mass, produced wet diarrhea-like frass (Fig 10G), ceased eating, began to change in color (melanization), as well as underwent a reduction in internal fluid pressure which led to deflation and overall lack of internal structure (Fig 10A-D). Dead larvae turned a pale green color, reflecting the color of the PAO1 bacterial colonies (Figure 10D). This coloration is a stark contrast to the vivid turquoise and blue of an uninfected individual (Fig 10E). Symptomatic hornworms were scored dead once they no longer reacted to external stimulus, such as prodding from forceps.

**Fig 10.**
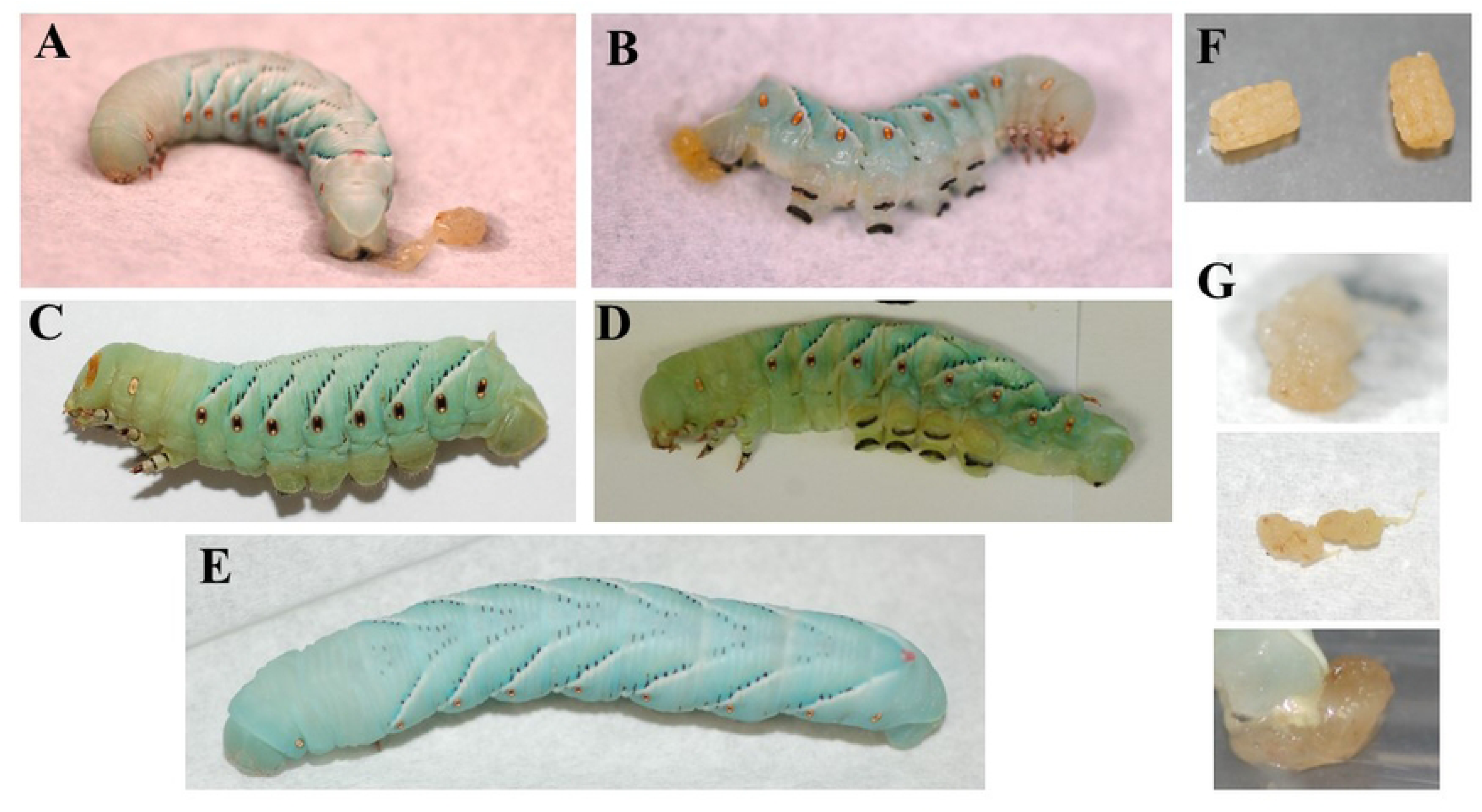
Symptomatic hornworm larvae compared to a healthy hornworm and frass. Hornworms that are exhibiting symptoms such as losing turgor pressure, producing wet, diarrhea-like frass, and have changed colors are shown in panels A, B, and C – as compared to the health hornworm shown in panel E. A dead hornworm, deflated and pale green is shown in panel D. Frass produced by a health hornworm is shown in F, as compared to the 3 examples in G of unhealthy, symptomatic hornworm frass.

## Discussion

In this study, we explored the utility of *Manduca sexta* larvae as an invertebrate model for studying bacterial pathogenesis and antibiotic efficacy. Although waxworms (*Galleria mellonella*) are widely used due to their simplicity and low cost, their use is often limited to survival assays. By contrast, the larger size and physiological complexity of *M. sexta* larvae allows for simultaneous measurement of survival, relative growth, and spaciotemporal bacterial burden, providing a more comprehensive picture of host-pathogen interactions.

Our results demonstrate that *M. sexta* larvae are susceptible to systemic infections from *P. aeruginosa* PAO1 and exhibit dose-dependent mortality and morbidity. High doses of bacteria were almost invariably lethal, whereas some words survived at lower doses but nonetheless experienced growth suppression. Thus, larval mass proved to be a sensitive indicator of sublethal infection effects within the five-to-six-day observation period. Infected larvae exhibited a range of disease symptoms, including an actual decline in mass, melanization, deflation, and eventual death, mirroring signs of infection observed in mammalian systems [67, 68]; however, these alternative phenotypes are not as easily scored as mass and survival. The dose-dependent mortality observed parallels findings from other model systems as well [66, 69–74].

Antibiotic intervention (gentamicin and cefepime) significantly improved growth and survival following an infection of live *P. aeruginosa*. In contrast, while antibiotic treatment markedly improved survival in larvae exposed to heat-killed PAO1, it did not significantly enhance growth among surviving individuals. This decoupling of mortality and growth responses suggests that distinct mechanisms underlie lethal versus sublethal effects of bacterial exposure [75, 76]. Survival alone therefore underestimates the physiological cost of the infection, and the ability to quantify growth trajectories reveals treatment effects that would be obscured in smaller invertebrate models, such as the waxworms [77–81].

Sequential hemolymph sampling revealed pronounced inter-individual variability in infection dynamics. Some larvae experienced sustained bacterial expansion, accompanied by reduced growth and eventual mortality. Others progressively cleared the bacteria and maintained near-normal growth. This heterogeneity indicates that infection outcomes depend on individual differences in the host’s ability to control bacterial proliferation, rather than reflecting a uniform pathogenic process [82–85].

Whole-body burden assays further clarified these dynamics. Although total internal bacterial concentration remained relatively stable over time, bacteria were redistributed from the hemolymph into the fat body, gut, and carcass tissues as the infection progressed. Elevated bacterial counts in the frass suggest that replication may have been offset by active excretion, contributing to the absence of net bacterial accumulation [86–88]. As might be expected, PAO1 replicated extensively in dead larvae, increasing substantially in the absence of the host’s immune system. This contrast indicates that bacterial proliferation is not intrinsically constrained by the larval tissue environment but is actively suppressed in living hosts [89–91]. High bacterial counts in the frass further raise the possibility of active replication and excretion, despite the minimal overall bacterial expansion in host tissues. These findings reflect complex host-pathogen dynamics involving immune responses, tissue invasion and clearance, and highlight the limitations of hemolymph-only sampling. The large size of *M. sexta* larvae uniquely enables whole organism and tissue specific dissection, providing critical spatial resolution not achievable in smaller invertebrate models.

Injection of heat-killed bacteria and filtered supernatant caused both mortality and morbidity, indicating that bacterial components such as toxins and/or immune-activating molecules can contribute heavily to host damage independent of a live infection [92–94]. The observation that heat-killed PAO1 caused greater mortality than antibiotic treated live infection underscores the contribution of bacterial products to pathogenesis and highlights the sensitivity of this system to both replicative and non-replicative virulence factors [92, 95–99].

Overall, this study highlights *M. sexta* as a useful invertebrate model for infectious disease research. Its large size enables simultaneous assessment of survival, growth, tissue-specific bacterial burden, and responses to inactivated bacterial components, with obvious advantages to understanding many dimensions of infections. While variability among individuals is substantial, this heterogeneity mirrors the complexity observed in vertebrate hosts and enhances the translational relevance of the model [100–102]. *M. sexta* therefore offers a bridge between simpler and smaller invertebrate systems, and more complex vertebrate models.

## Acknowledgements

The authors would like to thank Dr. Holly Wichman, and all of those involved with the Wichman-Miller Laboratory group at the University of Idaho for their advice, support, and expertise.

## Supporting Information

### Appendix. Statistical analysis tables

**Table A1. Dose-dependent mortality hazard ratios (Fig 1).** Hazard Ratios relative to the Controls for each of the PAO1 doses.

**Table A2. Dose-dependent growth statistics (Fig 2).** Day 4 pairwise comparisons of relative growth among surviving larvae.

**Table A3. Dose-dependent growth statistics (Fig 2) Tukey-adjusted pairwise contrasts**. Pairwise comparisons of treatment groups for larval relative growth at the experimental endpoint, based on estimated marginal means from a linear model including treatment group as a fixed effect. Reported values are Tukey-adjusted contrasts (estimate ± SE), with associated t-ratios and adjusted p-values (α = 0.05). Positive estimates indicate greater relative growth in the first-listed groups.

**Table A4. Individual variation in Growth (Fig 3).** The standard deviation (heterogeneity) of surviving larvae on day 4 post injection for each group and increasing bacterial dose.

**Table A5. Antibiotic treatment relative growth and controls (Fig 8).** Pairwise comparisons of treatment groups for larval relative growth at the Experimental endpoint, based on estimated marginal means from a linear model including treatment group as a fixed effect. Reported values are Tukey-adjusted contrasts (estimate ± SE), with associated t-ratios and adjusted p-values (α = 0.05). Positive estimates indicate greater relative growth in the first-listed groups.

**Table 6A. Survival of different treatments: live cells and PAO1 byproducts (Fig 9).** Hazard ratios (HR) are shown relative to live PAO1 infection. Values less than 1 indicate reduced mortality risk relative to PAO1. All treatments significantly reduced mortality risk compared to live infection. Antibiotic treatment, particularly when combined with heat-killed bacteria, provided strongest protection. Proportional hazards assumption was satisfied for all comparisons.

**Table 7A. Relative growth of different treatments: live cells and PAO1 byproducts (Fig 9).** Pairwise comparisons of treatment groups for larval relative growth at the experimental endpoint, based on estimated marginal means from a linear model including treatment group as a fixed effect. Reported values are Tukey-adjusted contrasts (estimate ± SE), with associated t-ratios and adjusted p-values (α = 0.05). Positive estimates indicate greater relative growth in the first-listed groups.

**S1 Dataset. Raw weights of larvae for treatment groups.** This dataset shows raw weights (in grams) over time for each group of larvae mentioned in this manuscript. The relative weights/growth are shown in figures, and the raw weights are shown in this dataset.

---

**Fig S1.**
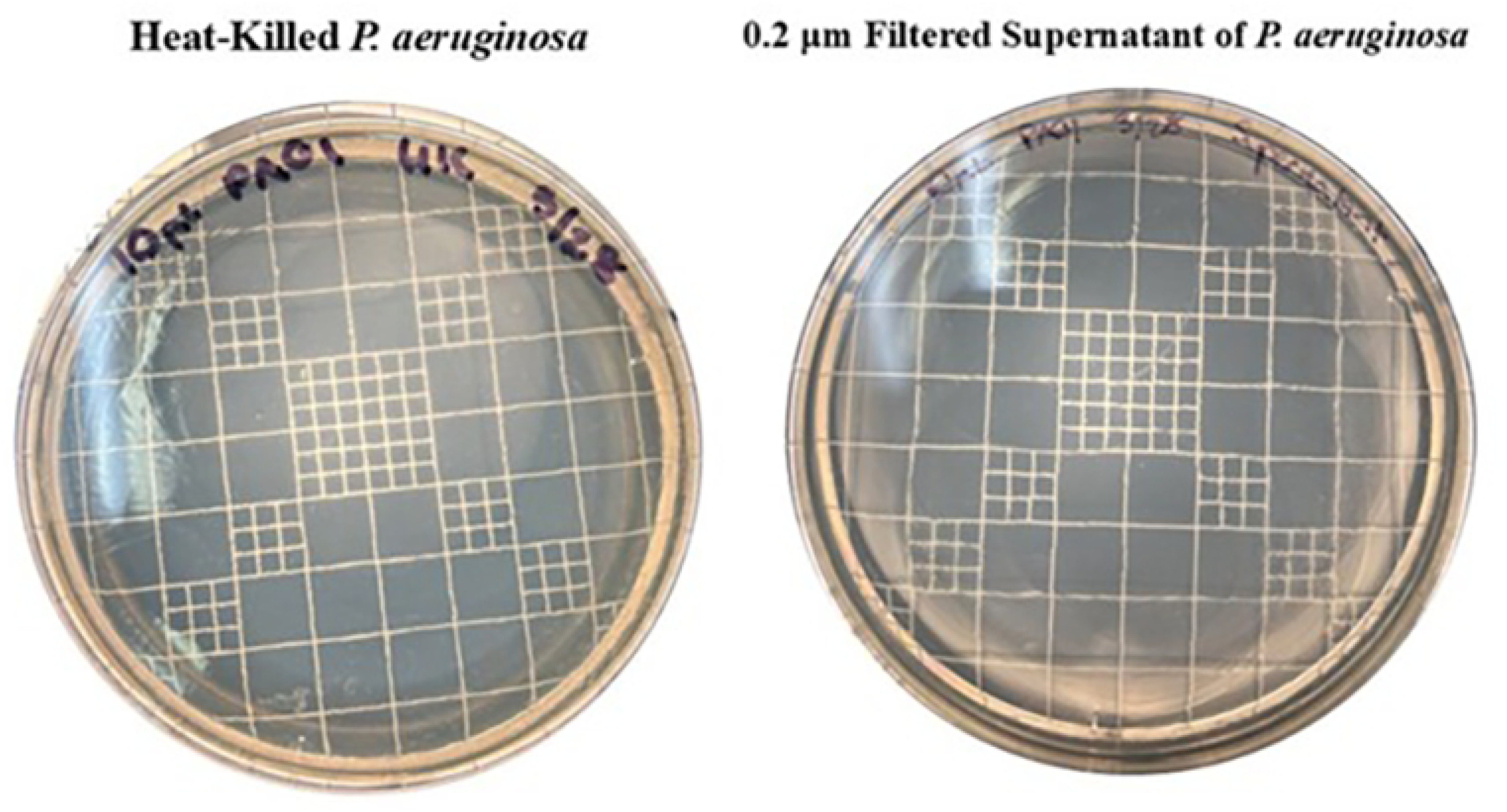
Heat-killed and filtered supernatant on LB plates to determine presence of live colonies. 10 microliters of Heat-killed PAO1 and 10 microliters of Filtered Supernatant PAO1 on LB plates after incubation at 37 °C overnight. No bacterial colonies are present, indicating that there are no live, active bacterial cells that are replicating.

**Fig S2.**
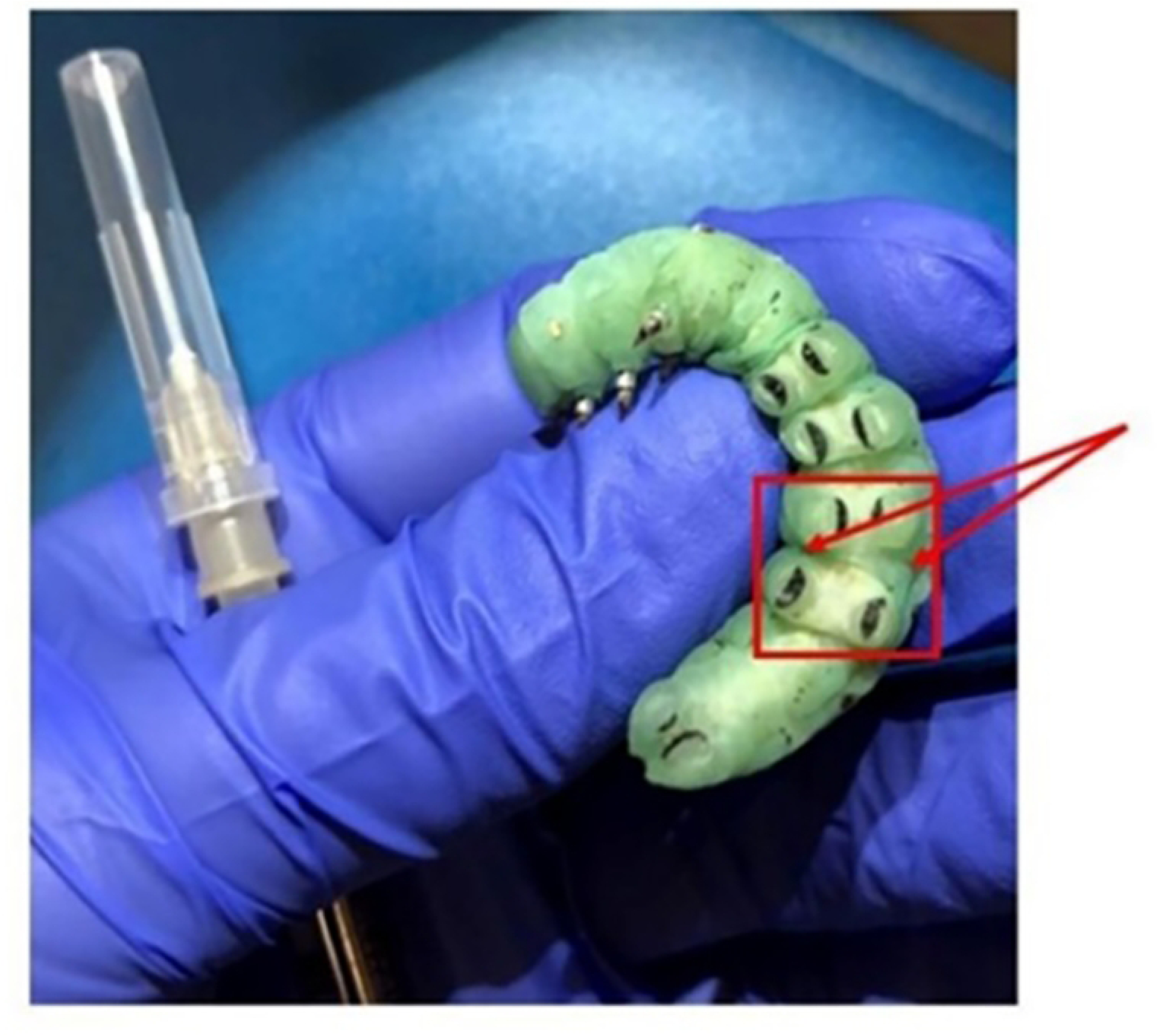
Location for injections on fifth instar larvae. The injections were done with a 30-guage needle in the crease between the third and the fourth prolegs.

**Fig S3.**
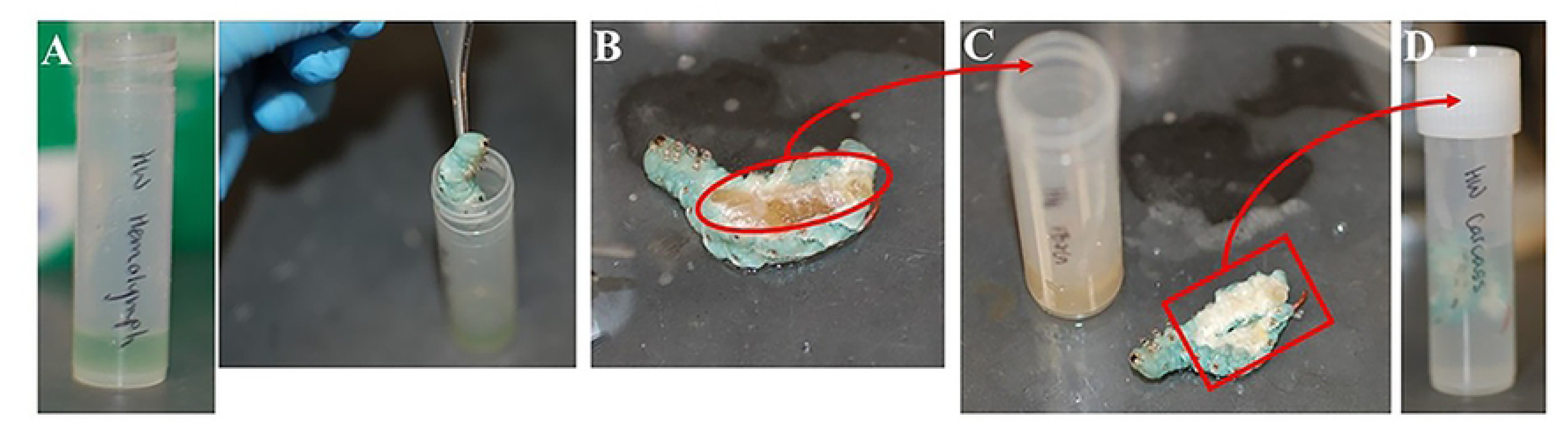
Dissected tissues of hornworm larvae. A) Hemolymph collection. B) Fat body and gut tract dissection and collection. C) Carcass collection is what remains from removing the fat body and gut tract, as well as the head. D) Tube with the carcass suspended in PBS.

**Fig S4.**
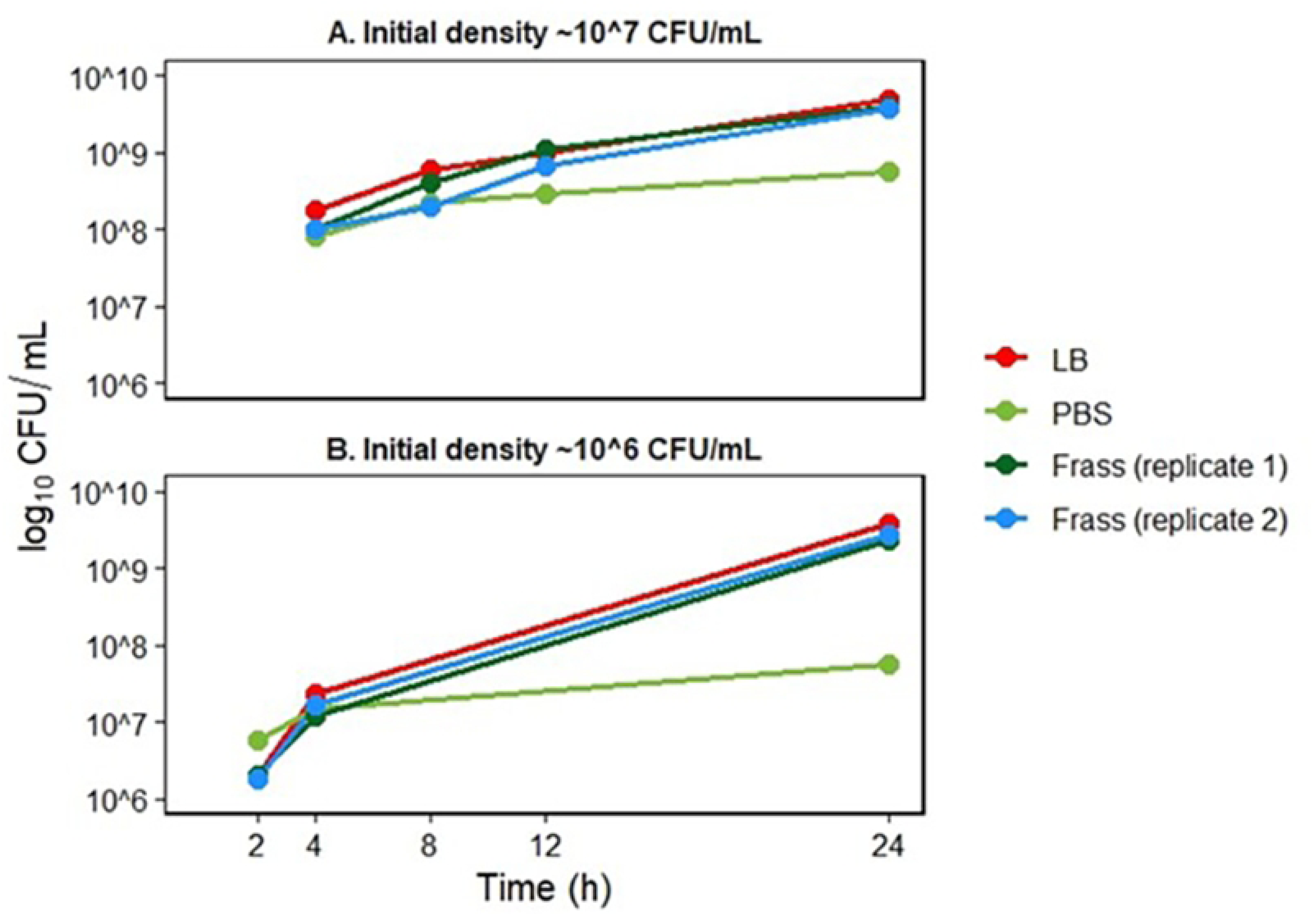
PAO1 growth in LB, PBS, and hornworm frass. Bacterial growth was measured in LB, PBS, and frass across two assays initiated at different starting densities: A) ∼10^7^ CFU/mL and B) ∼10^6^ CFU/mL. In both assays, PAO1 increased substantially in frass and reached densities more similar to those observed in LB than in PBS. Frass replicates are shown separately.

